# *In vitro* to *in vivo* pharmacokinetic translation guidance

**DOI:** 10.1101/2022.09.26.509470

**Authors:** Urban Fagerholm

## Abstract

**Background:** Pharmacokinetics (PK), exposure profiles and doses of candidate drugs in man are commonly predicted using data produced using various *in vitro* methods, such as hepatocytes (for intrinsic metabolic clearance (CL_int_)), plasma (for unbound fraction (f_u_)), Caco-2 (measuring apparent permeability (P_app_) for prediction of *in vivo* fraction absorbed (f_a_)) and plasma water and buffers (measuring solubility (S) for prediction of *in vivo* fraction dissolved (f_diss_)). For best possible predictions it is required that the clinical relevance of *in vitro* data is understood (*in vitro-in vivo* relationships) and that uncertainties have been investigated and considered.

**Methods:** The aim was to investigate *in vitro-in vivo* relationships for CL_int_, P_app_ *vs* f_a_ and S *vs* f_diss_ and interlaboratory variability for f_u_, describe the clinical significance and uncertainties at certain levels of *in vitro* CL_int_, f_u_, P_app_ and S, and (based on the findings) develop a general *in vitro-in vivo* translation guide.

**Results and Conclusion:** It was possible to finf data for describing how *in vivo* CL_int_, f_a_ and f_diss_ distribute and varies at different levels of *in vitro* CL_int_, P_app_ and S and how f_u_ varies between laboratories and methods at different f_u_-levels. It is apparent that there are considerable interlaboratory variabilities for CL_int_, f_u_ and P_app_: corresponding to up to 2500-, 700- and 35-fold variability for CL_int_, f_u_ and f_a_, respectively. Apparently, S is a poor predictor of f_diss_. Proposed S-thresholds do not seem clinically useful (overestimated). It does not seem appropriate to define *in vitro* CL_int_ of 0.5-2 µL/min/10^6^ cells as good metabolic stability (rather moderate to moderately high). Results shown for CL_int_, P_app_ and f_u_ are applicable as general guidelines when internal standard values for reference compounds are unavailable.

## Introduction

Pharmacokinetics (PK), exposure profiles and doses of candidate drugs in man are commonly predicted using data produced using various *in vitro* methods, such as human hepatocytes (for measuring intrinsic metabolic clearance (CL_int_) and predicting ditto *in vivo* in man), plasma (for measuring unbound fraction (f_u_) and assuming that it resembles the *in vivo* situation), Caco-2 (measuring apparent permeability (P_app_) for prediction of *in vivo* fraction absorbed (f_a_)) and plasma water and buffers (measuring solubility (S) for prediction of *in vivo* fraction dissolved (f_diss_)).

*In vitro* PK-data are laboratory and method dependent, which means that each laboratory aiming for optimal predictions needs to establish their own *in vitro-in vivo* relationships using data for reference compounds of various characteristics and with different PK-properties. Apparently, this is not always the case. It has also been shown that sometimes data for compounds with non-quantifiable PK-properties and outliers are avoided or excluded and that favorable data are selected (which leads to excessively positive results and strong correlations) (1).

For best possible predictions it is also required that the clinical relevance of *in vitro* data is understood (*in vitro-in vivo* relationships) and that uncertainties and outliers have been investigated and considered.

The aim of the study was to investigate *in vitro-in vivo* relationships for CL_int_, P_app_ *vs* f_a_ and S *vs* f_diss_ and interlaboratory variability for f_u_, describe the clinical significance and uncertainties at certain levels of *in vitro* CL_int_, f_u_, P_app_ and S, and, based on the findings, develop a general *in vitro-in vivo* translation guide.

## Methods

### Data selection

The literature was searched for data for human hepatocyte CL_int_, f_u_, Caco-2 P_app_ and water or buffer S, and for corresponding clinical CL_int_, f_u_, f_a_ and f_diss_. Data were taken from the following references: human hepatocyte CL_int_ (2–8), f_u_ (9,10), Caco-2 P_app_ (11–21), S (12 and collection of data proprietary of Prosilico AB), *in vivo* CL_int_ (22), f_a_ (12,22) and f_diss_ (data proprietary of Prosilico AB).

### *In vitro-in vivo* relationships and data levels

*In vivo* estimates and ranges at certain *in vitro* estimate levels were investigated. The selected *in vitro* levels were <LOQ (<limit of quantification (ca <0.5-2 µL/min/10^6^ cells) or non-measurable), 2 (1–3) µL/min/10^6^ cells (approximate LOQ for the conventional hepatocyte assay) and 10 (7–13) µL/min/10^6^ cells for CL_int_, 0.005 (0.004-0.006), 0.05 (0.04-0.06) and 0.50 (0.40-0.60) for f_u_, 0.1, 1 and 10 • 10^−6^ cm/s for Caco-2 P_app_, and 1 (0.7-1.5), 10 (7–15) and 100 (70-150) µM for S (suggested minimum S required for minimum acceptable f_a_ of high permeability-compounds at oral doses of 7, 70 and 700 mg, respectively (23)).

## Results

### CL_int_

Figure 1 shows *in vivo* CL_int_ at <LOQ (n=59), 2 µL/min/10^6^ cells (n=36) and 10 µL/min/10^6^ cells (n=18). Median *in vivo* CL_int_ at these *in vitro* levels are ca 400, ca 1200 and ca 10500 mL/min, respectively. Corresponding minimum and maximum levels are ca 30 and >15000 mL/min (>470-fold range), ca 5 and ca 15000 mL/min (2500-fold range), and ca 600 and ca 700000 mL/min) (1200-fold range), respectively. The median *in vivo* CL_int_ for marketed drugs up to year 2010 is 738 mL/min (22), which is half of the liver blood flow rate and higher than for 50 (44 %) of the 113 compounds. The 70 % confidence interval (CI) at <LOQ, 2 µL/min/10^6^ cells and 10 µL/min/10^6^ cells are 225-ca 4000 (18-fold), 318-3750 (12-fold) and 1575-56250 (36-fold) mL/min, respectively.

**Figure 1.**
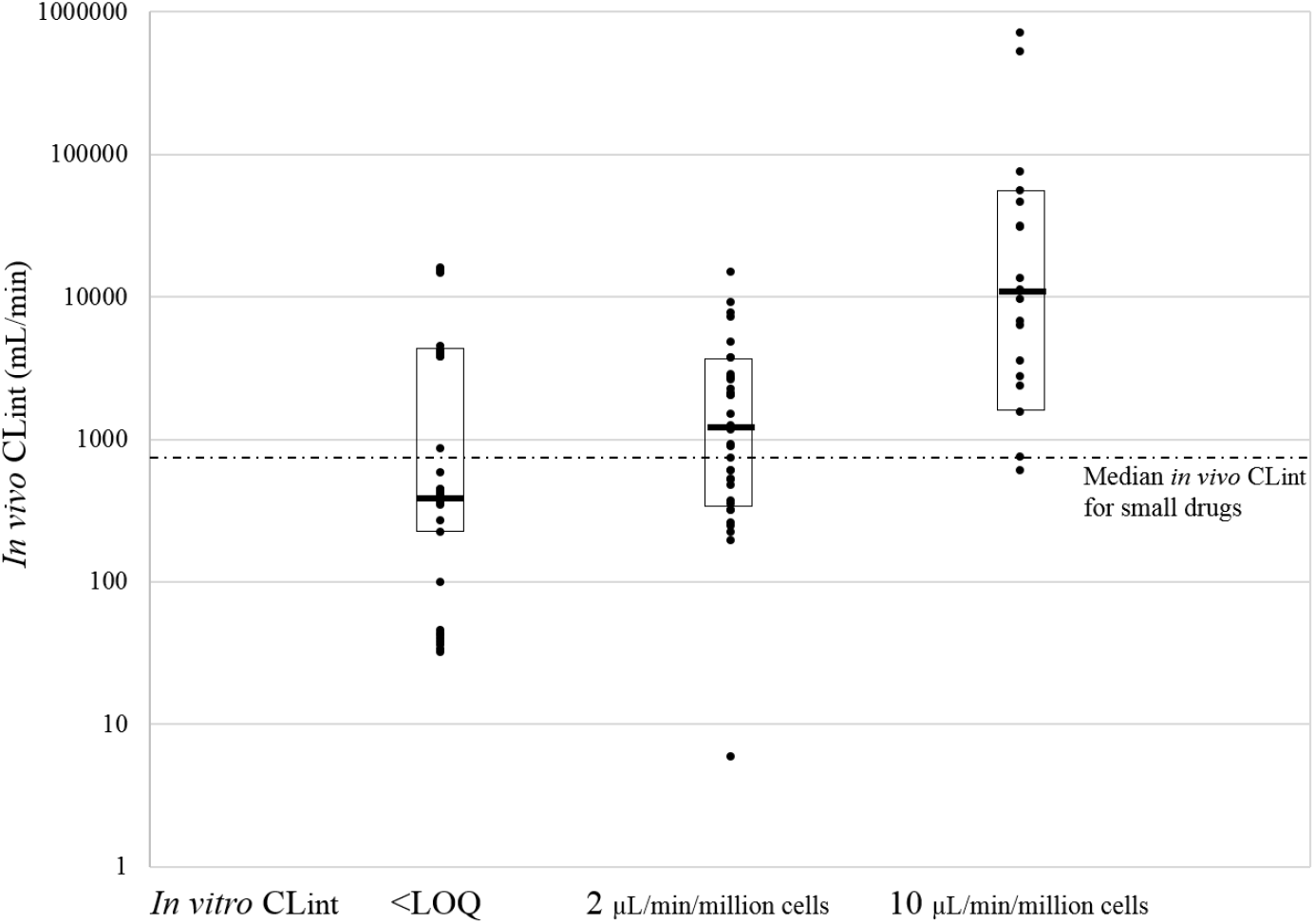
Individual *in vivo* CL_int_ at <LOQ (n=59), 2 µL/min/10^6^ cells (n=36) and 10 µL/min/10^6^ cells (n=18), median *in vivo* CL_int_ for marketed small drugs (dotted line), and median (-) and 70 % CIs (boxes) for each group.

### f_u_

In Figure 2, reported f_u_ at f_u_-levels 0.005 (n=20), 0.05 (n=31) and 0.50 (n=6) are shown. For a compound with a f_u_ of 0.005 in one laboratory other laboratories may (based on results from the collected data) produce values of somewhere between ca 0.0001 and 0.07 (700-fold range) for that compound (Figure 2). Corresponding ranges for compounds with measured f_u_ of 0.005 and 0.50 in one laboratory are ca 0.001 to 0.50 (500-fold range) and 0.06 to 0.97 (16-fold range), respectively. Corresponding 70 % CI are 0.003-0.037 (12-fold), 0.01-0.068 (7-fold) and 0.14-0.88 (6.3-fold), respectively.

**Figure 2.**
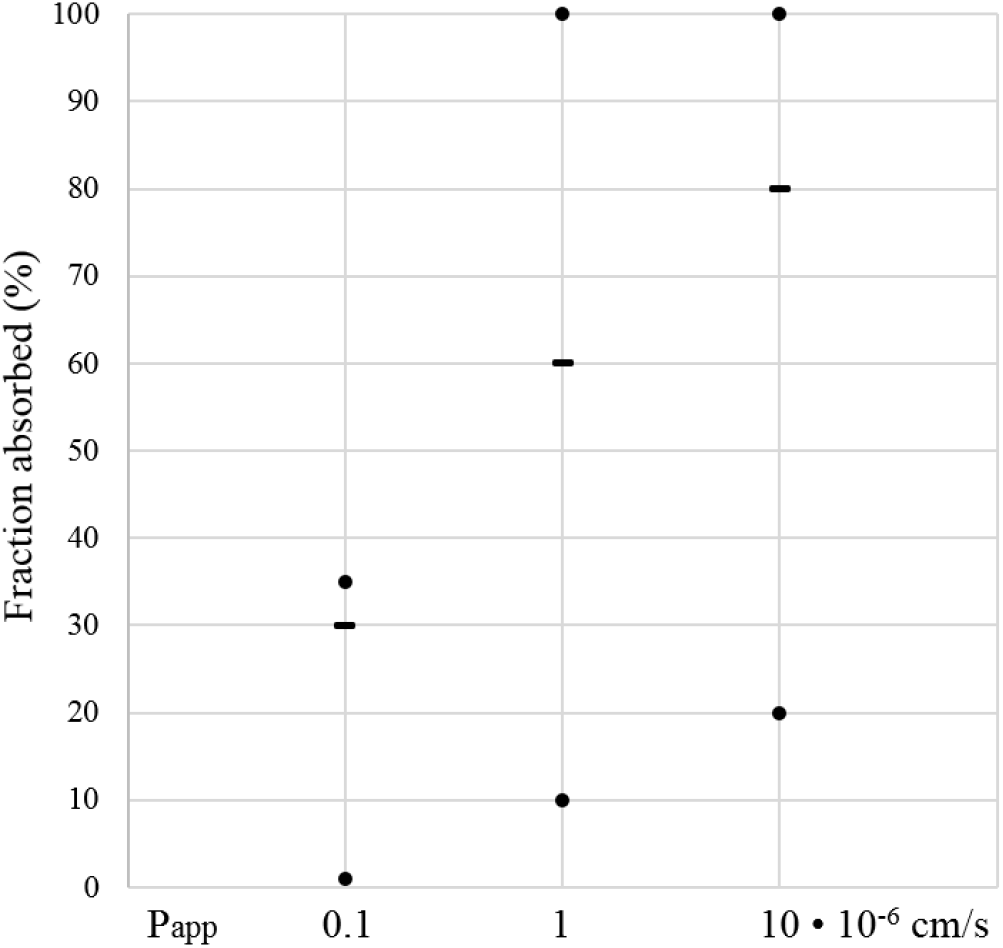
Reported f_u_ and estimated 70 % CIs (boxes) at f_u_-levels 0.005 (n=20), 0.05 (n=31) and 0.50 (n=6).

### P_app_

Pham-The et al. (12) established a typical sigmoidal relationship between log Caco-2 P_app_ and f_a_ (average of data from various sources; n=282) and found that for Caco-2 P_app_ 0.1, 1 and 10 • 10^−6^ cm/s the f_a_ averages ca 30, ca 60 and ca 80 %, respectively (Figure 3). Corresponding f_a_-ranges are ca 1-35 % (35-fold), ca 10 (intrapolated)-100 % (10-fold) and ca 20-100 % (5-fold), respectively. At a P_app_-level as for metoprolol (with f_a_=97 %; average P_app_=18 • 10^−6^ cm/s) f_a_ ranges between ca 25 and 100 %.

**Figure 3.**
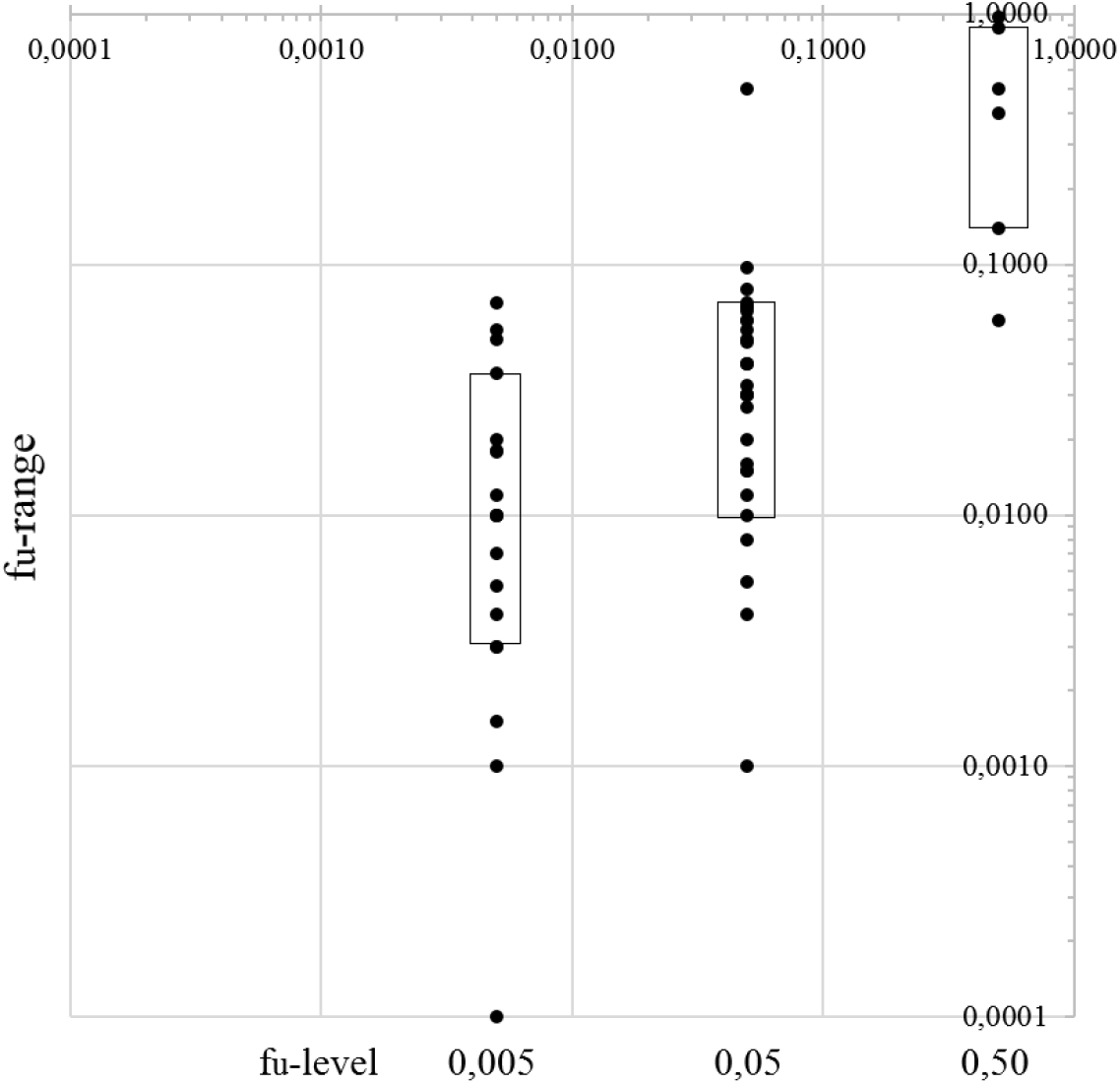
Minimum (•), median (-) and maximum (•) fraction absorbed (f_a_) for compounds with Caco-2 P_app_ 0.1, 1 and 10 • 10^−6^ cm/s (data taken from reference 12).

Several groups have found that Caco-2 P_app_ of ca ≥10 • 10^−6^ cm/s corresponds to f_a_≥80-90 %, but there are also reports where lower (ca 2-3 • 10^−6^ cm/s) or higher (ca 30 • 10^−6^ cm/s) Caco-2 P_app_ at and above this f_a_-limit (12, 19–21). A similar trend is apparent for Caco-2 P_app_ corresponding to a f_a_ of 50 %.

### S

Individual and median (-) maximum *in vivo* dissolution potential (f_diss_) for compounds with *in vitro* S of 1 (n=8), 10 (n=3) and 100 (n=7) µM are shown in Figure 4. All compounds in this S-range, except for the high oral dose-compounds atavaquone (S=1.2 µM; f_diss_=6 %; 500 mg oral dose), have a f_diss_ of ≥90 %. Median f_diss_ for the S-levels 1, 10 and 100 µM are 98, 100 and 100 %, respectively.

**Figure 4.**
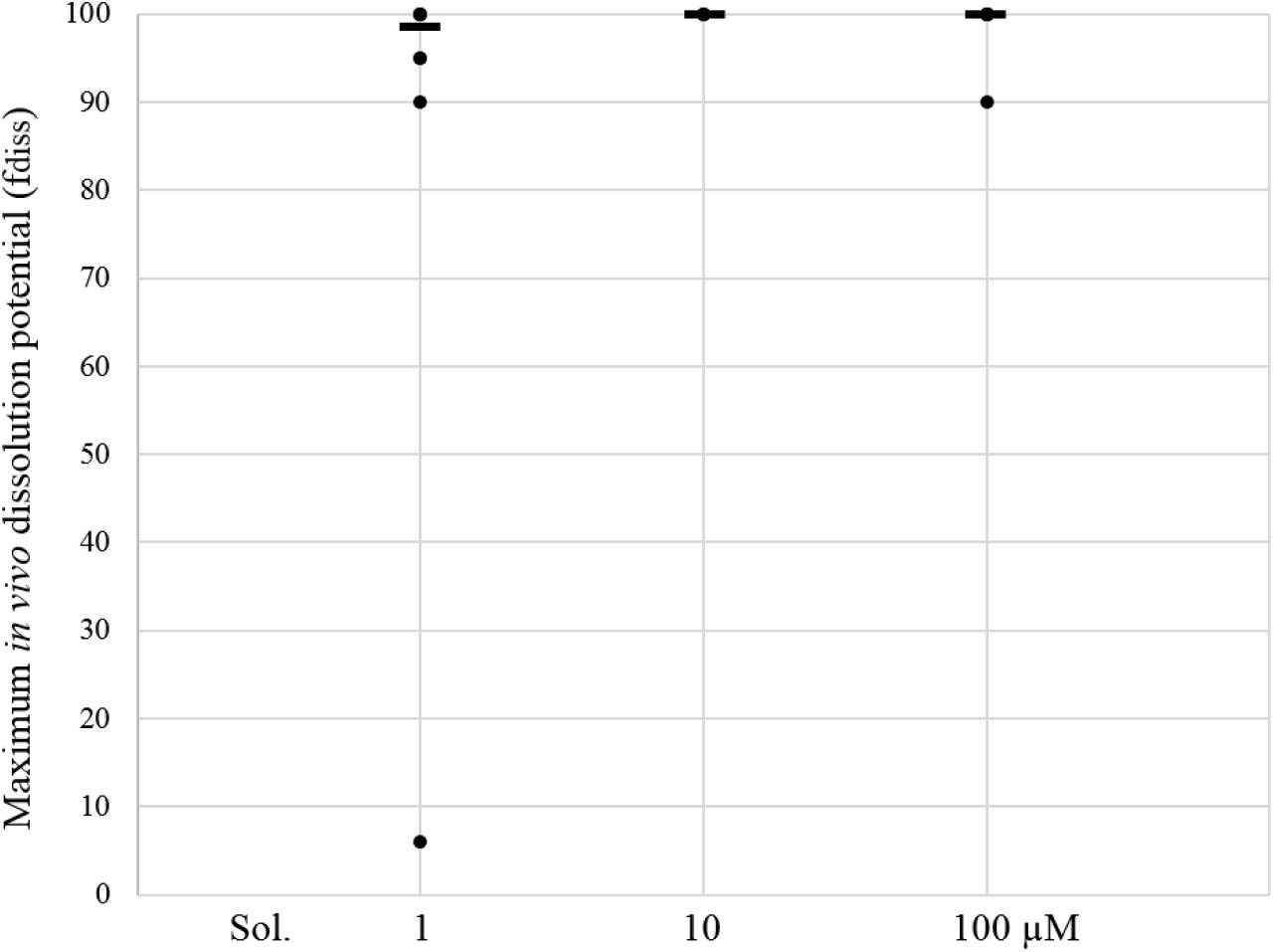
Individual (•) and median (-) maximum *in vivo* dissolution potential (f_diss_) for compounds with *in vitro* aqueous solubilities (S) of 1 (n=8), 10 (n=3) and 100 (n=7) µM.

## Discussion

It was possible to find data for describing how *in vivo* CL_int_, f_a_ and f_diss_ distribute and varies at different levels of *in* vitro CL_int_, P_app_ and S and how f_u_ varies between laboratories and methods at different f_u_-levels.

It is apparent that there are considerable variabilities for data produced and collected at different laboratories: up to 2500-, 700- and 35-fold for CL_int_, f_u_ and f_a_, respectively. We have found that interlaboratory variability is on average approximately half of that between laboratories (data on file).

Results shown for CL_int_, P_app_ and f_u_ are general, and should therefore, be used as general guidelines when internal standard values for reference compounds are unavailable.

Apparently, S is a poor predictor of f_diss_ at S≥1 µM. One explanation to this is that f_diss_-is dose-dependent. Another is that they are very different constructs. Results indicates that limits for S should be lower than previously proposed and that dose level needs to be considered.

Results for CL_int_ demonstrate that *in vitro*-estimates at LOQ (ca 0.5-2 µL/min/10^6^ cells) correspond to a comparably high *in vivo* CL_int_ (near the median *in vivo* CL_int_ for small drugs) and that compounds with *in vitro* CL_int_≤2 µL/min/10^6^ cells can have an *in vivo* CL_int_ of up to 15000 mL/min (10 times higher than the lover blood flow rate). Thus, it does not seem appropriate to define *in vitro* CL_int_ of 0.5-2 µL/min/10^6^ cells as good metabolic stability (rather moderate to moderately high).

For gemfibrozil, 51- and 18-fold interlaboratory variabilities have been shown for *in vitro* CL_int_ and f_u_ (3,9). The implication of this is that two laboratories may show almost 1000-fold different Cl_int •_ f_u_. This result further shows the requirement for the use of reference compounds and internal standards when predicting the human clinical PK, but also the possibility to select favorable data, risk to make wrong decisions (compound selection and stop-go) and overdose individuals in early clinical trials with new drug candidates.

## Conclusion

*In vitro-in vivo* relationships for CL_int_, P_app_ *vs* f_a_ and S *vs* f_diss_ and interlaboratory variability for f_u_ were established, the clinical significance and uncertainties at certain levels of *in vitro* CL_int_, f_u_, P_app_ and S were described and a general *in vitro-in vivo* translation guide was developed.

